# The transcription factor STAT5 binds to distinct super-enhancer sites and controls *Lrrc32* expression in a prominent autoimmune and allergic disease risk locus

**DOI:** 10.1101/2020.06.13.150177

**Authors:** Lothar Hennighausen, Hye Kyung Lee

**Affiliations:** Laboratory of Genetics and Physiology, National Institute of Diabetes and Digestive and Kidney Diseases, US National Institutes of Health, Bethesda, Maryland 20892, USA

## Abstract

Genetic variants associated with diseases are enriched in genomic sequences linked to regulatory regions, such as enhancers, super-enhancers and possibly repressors, that control nearby and distant genes. A known allergic and autoimmune risk locus at chromosome 11q13.5^1,2^ is associated with the *LRRC32* gene, which encodes GARP, a protein critical for TGF-β delivery^3^. This region coincides with a candidate enhancer that was predicted by the presence of activating chromatin marks and contains a polymorphism significantly associated with GARP expression on CD4^+^CD127^-^CD25^+^ T_reg_ cells^4^. In the mouse, binding of the cytokine-induced transcription factor STAT5 was detected at two sites within the expansive candidate enhancer region and a 2.3 kb deletion resulted in reduced *Lrrc32* expression^4^. However, a clear definition of the enhancer units controlled by STAT5 and a functional understanding of STAT5 in the regulation of *Lrrc32* are needed. Here we use high-resolution ChIP-seq and identify three STAT5 binding sites within the *Lrrc32* super-enhancer, one shared between T_reg_ cells and mammary epithelium and one specific to each respective cell type. Using mice that express only 10% of normal STAT5 levels we demonstrate the defining contribution of STAT5 in the activation of the *Lrrc32* super-enhancer.

## Introduction

Genome-wide approaches, such as whole genome sequencing and ChIP-seq based technologies, have provided researchers with unique handles to understand molecular underpinnings of gene regulation and the nature of disease susceptibility loci. It turns out that disease susceptibility loci are frequently found in chromosomal regions that harbor genetic regulatory elements, such as enhancers and superenhancers, and mutations in transcription factor motifs could alter the binding affinities of regulatory proteins^5^. Accurately defining the linear dimensions of enhancers and super-enhancers, the exact positions of transcription factors on the DNA, and the intricacy of the regulatory complex requires high-resolution ChIP-seq technologies.

Nasrallah and colleagues^4^ have investigated the molecular structure and biology of a candidate enhancer in a prominent autoimmune and allergic disease risk locus at chromosome 11q13.5. This enhancer was predicted by the presence of the activating chromatin mark K27-acetylated histone H3 (H3K27ac) in CD4^+^CD127^-^CD25^+^ regulatory T cells and other primary lymphocyte lineages. This region is approximately 70 kb downstream of the LRRC32 gene that encodes GARP, a protein critical for the function of Foxp3^+^ CD4^+^ regulatory T cells, also known as T_reg_ cells^6^. Deletion of a 2.3kb enhancer fragment from the syntenic mouse region resulted in reduced GARP levels in T_reg_ cells and the inability to control colitis^4^. A putative binding motif for the cytokine-inducible Signal Transducer and Activator of Transcription (STAT) 5 was identified by DNA sequences and a ChIP-qPCR assay revealed STAT5B binding, suggesting its contribution in the activation of *Lrrc32*. However, the proposed STAT5 motif (AGGAAATA) is distinctly different from the canonical palindromic GAS motif (TTCNNNGAA). Since ChIP-qPCR monitors transcription factor binding over several hundred nucleotides it is insufficiently precise to accurately pinpoint transcription factor (TF) binding sites. It is therefore possible that STAT5 binds to sequences other than those proposed^4^, or piggybacks on unrelated TFs or even that additional non-STAT TFs might control this enhancer.

## Results and Discussion

To clearly identify and characterize the candidate enhancer linked to the *Lrrc32* gene, we turned to high-precision ChIP-seq data from human and mouse and experimental mouse genetics. The transcription factor STAT5A and STAT5B (referred to as STAT5) have non-redundant and essential roles in the physiology of T_reg_ cells and mammary epithelial cells^7–9^. Here, by integrating high-quality ChIP-seq, RNA-seq and genetic data from T_reg_ cells^10–13^ and mammary tissue^14,15^, we define the structure of the *Lrrc32* enhancer, accurately identify the binding positions of STAT5 and other TFs, and elucidate the significance of STAT5 in regulating *Lrrc32* expression.

First, we analyzed ChIP-seq data from human T cells^13^ (Fig. 1). The H3K27ac pattern predicted the presence of several candidate enhancer regions between the 3’ ends of the *Lrrc32* and *Emsy* genes. The most prominent one was ~70 kb 3’ of the *Lrrc32* gene (Fig. 1a). This region also featured two sites, E1 and E2, bound by STAT5. Sequences around site E1 are conserved between human and mouse (Fig. 1b) and a STAT5 binding site had been predicted (green highlight in Fig. 1b), next to the SNV that was linked to reduced GARP levels^4^. However, STAT5 does not recognize this site and it rather binds to a GAS (TTCNNNGAA) motif at a distance of 70 bp (yellow highlight in Fig. 1b). STAT5 binding at site E2 also coincides with a GAS motif, the consensus binding motifs for all STAT members except STAT6.

**Fig. 1.**
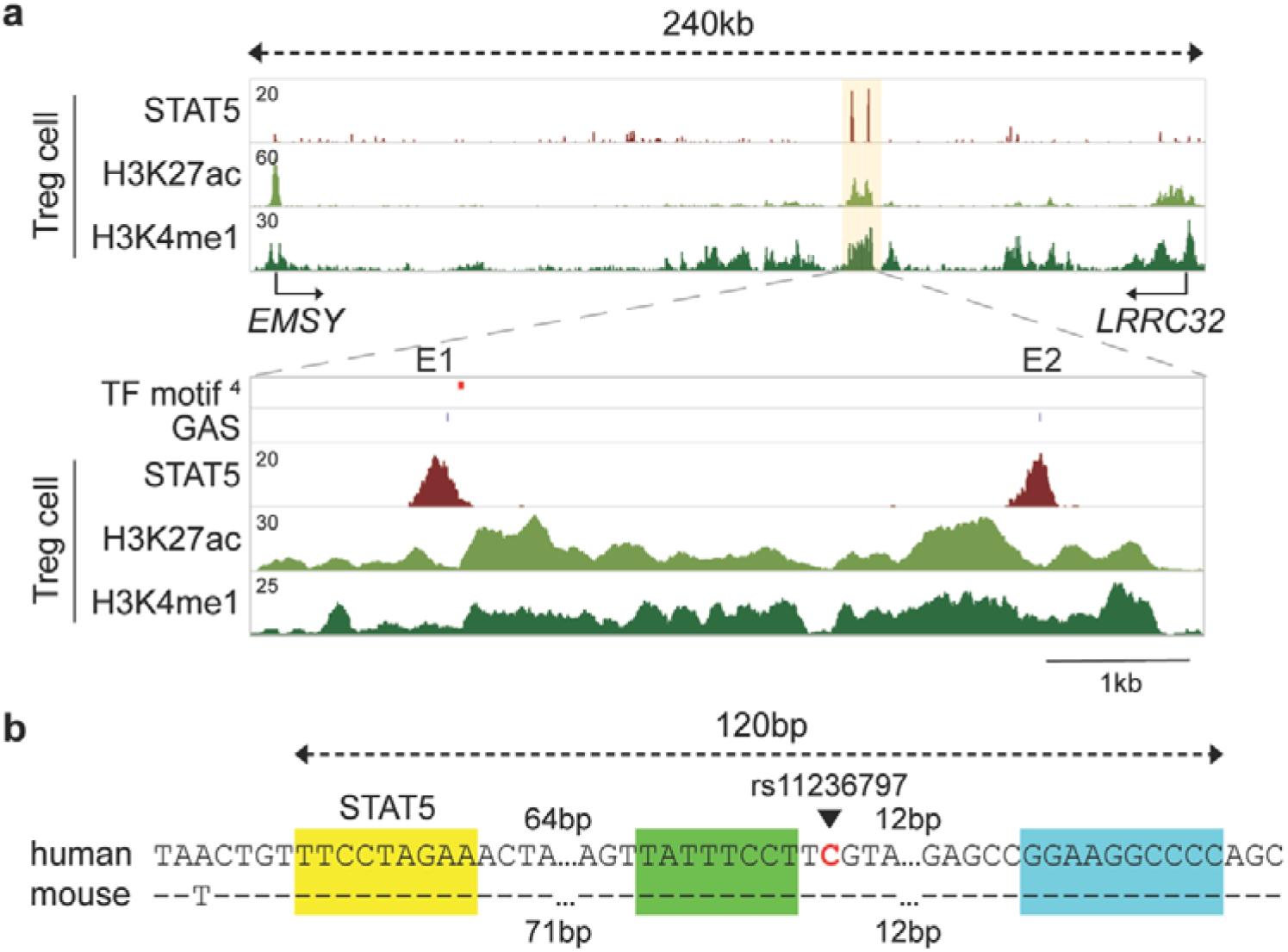
Conserved transcription factor binding motifs in the *LRRC32* enhancer region. **a,** ChIP-seq profiles showed STAT5 binding at the *LRRC32* locus and the candidate enhancers in human T_reg_ cells. The orange shades indicate candidate regulatory regions. The canonical GAS motifs are indicated by a blue marks (E1 and E2). The ‘STAT site’ identified by Nasrallah^4^ is indicated by a red mark. E1 and E2 are putative enhancers with strong STAT5 binding in human T_reg_ cells. **b,** Genomic sequence alignment (hg19 and mm10) shows the conserved canonical STAT5 binding motif (GAS) in yellow, the ‘STAT5 site’ identified by Nasrallah^4^ highlighted in green and the NF-κB binding motif (Nasrallah^4^) highlighted in turquois.

Next, we dug deeper and analyzed the structure and significance of these sites using ChIP-seq data from mouse T_reg_ cells^10–12^ and mouse genetics^10,15^. Like in human T cells, STAT5 binding at site E1 coincides with a GAS motif and is located approximately 90 bp apart from the site identified in the Nasrallah study^4^ (Figs. 1b and 2). The presence of H3K27ac marks and the H3K4me1 signature, which identifies genuine enhancers, support the regulatory nature of this region. Also, site E2 was present and coincides with all activating histone marks was discovered (Fig. 2a).

**Fig. 2.**
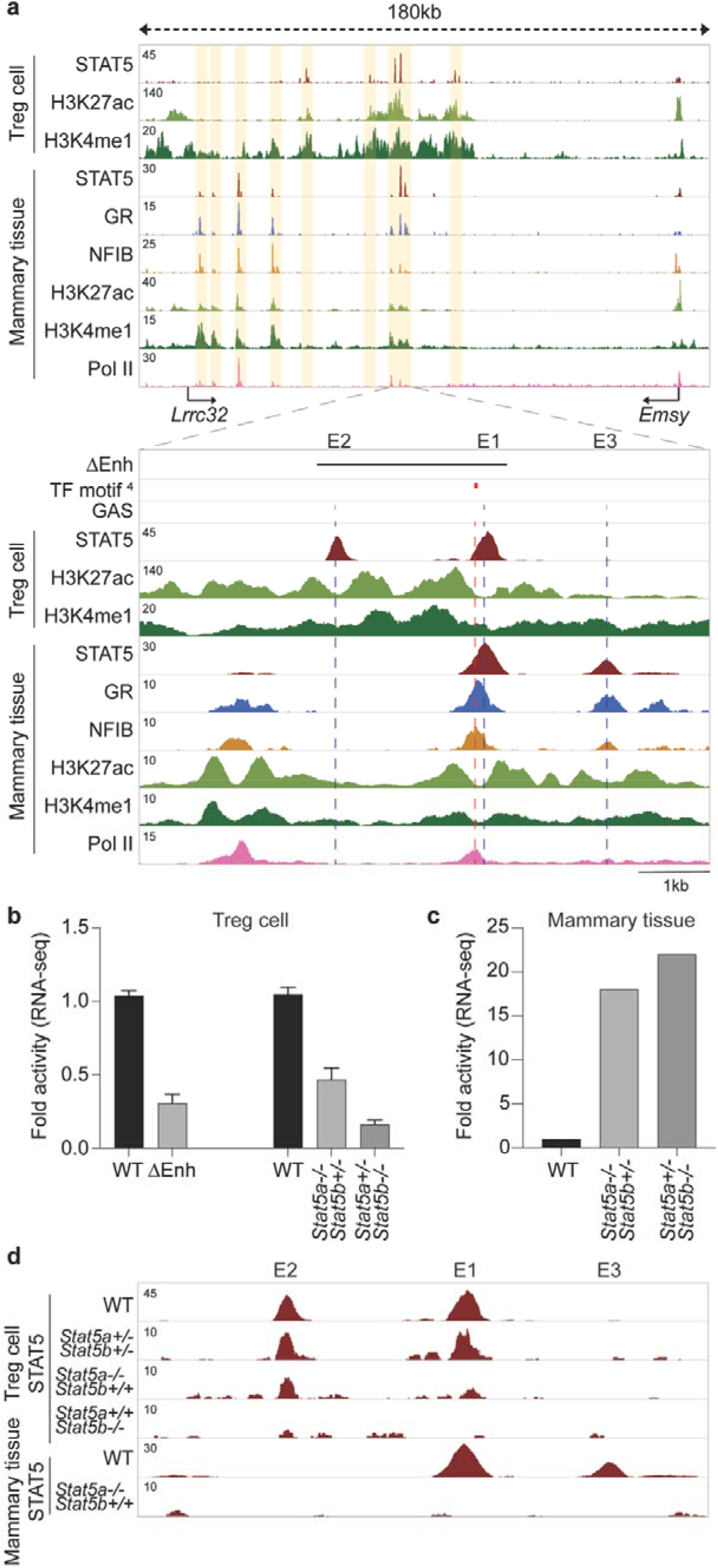
Establishment of putative *Lrrc32* enhancers in mammary epithelium and regulatory T (T_reg_) cells. **a,** ChIP-seq data for STAT5A (in mammary tissue) or panSTAT5 (in IL2-stimulated CD4+ T cells), GR, NFIB, Pol II and histone marks H3K27ac and H3K4me1, provide structural information of the locus between the *Lrrc32* and *Emsy* genes in lactating mammary tissue and activated T_reg_ cells. Solid arrows indicate the orientation of genes. The orange shades highlight candidate regulatory regions. The black bar indicates the deleted enhancer region (ΔEnh)^4^. The blue broken lines (E1, E2 and E3) highlight STAT5 binding associated with the canonical GAS motif. The red line depicts the ‘STAT site’ identified by Nasrallah^4^. E1, E2 and E3 are putative enhancers with strong STAT5 binding. **b,** RNA-seq data from Nasrallah^4^ and Villarino^10^ demonstrate the relative expression of *Lrrc32* gene in T_reg_ cells in the absence of the 2.3kb enhancer region and different *Stat5* alleles, respectively. **c,** *Lrrc32* mRNA levels in mouse mammary tissue of WT and *Stat5* mutant mice were measured by RNA-seq^15^. **d,** The putative enhancers were profiled using ChIP-seq data of WT and mutant tissue.

Since T_reg_ cells and mammary epithelium share their absolute dependence on STAT5^7,9^ we surmised that key enhancers might be shared between these two systems. Using high-quality ChIP-seq data from mouse mammary tissue^14,15^ we identified strong STAT5 binding at two juxtaposed sites (E1 and E3) (Fig. 2a). While site E1 is shared between T_reg_ cells and mammary tissue, site E3 appears to be mammary-specific. Also, site E3 does not coincide with the one identified in the Nasrallah study^4^. However, the TF site identified by Nasrallah et al^4^ is occupied by two other TF, the glucocorticoid receptor (GR) and nuclear factor I B (NFIB) (on E1 at bottom in Fig. 2a) suggesting these, rather than STAT5, are the occupiers of this enhancer region. The presence of the enhancer-distinctive patterns of H3K27ac and H3K4me1 marks, in particular the histone-depleted sequences surrounding the TF binding, and the presence of RNA Polymerase II (Pol II) support the regulatory significance of this region.

Vetting STAT5 and its role in activating the *Lrrc32* gene can be accomplished either through the deletion of the GAS motifs or the ablation of the *Stat5* genes. Nasrallah^4^ deleted a 2.3 kb fragment spanning enhancer sites E1 and E2 (Fig. 2a), which resulted in an approximately 65% reduction of *Lrrc32* expression in T_reg_ cells (Fig. 2b). Since this region is extensively covered by activating histone marks and harbors at least two distinct enhancers bound by several TFs, interpretation on the role of STAT5 in *Lrrc32* regulation cannot be definitive. We have now addressed the specific contribution of STAT5 in regulating *Lrrc32* gene expression in T_reg_ cells and mammary tissue through mutant mice that carry different copy numbers of STAT5 genes^15^. Since the combined deletion of *Stat5a* and *Stat5b* results in perinatal lethality^7,16^, we generated mice that lack three out of the four *Stat5* alleles and carry only a single *Stat5a* or *Stat5b* allele (Fig. 2b)^15^. In the presence of only one *Stat5a* allele, *Lrrc32* expression was reduced by ~85% in T_reg_ cells^10^ compared to wild type (WT) cells supporting the notion that STAT5 is a key component of the enhancer. Since STAT5B is the more abundant isoform in T_reg_ cells, the presence of one copy of *Stat5b* resulted in a ~55% reduction of *Lrrc32* expression. In contrast to T_reg_ cells, *Lrrc32* expression in mammary tissue increased 22-fold in the absence of three out of the four *Stat5* alleles (Figs. 2c and d), suggesting that STAT5 might be a context-specific repressor. Using ChIP-seq from our allele-specific mutants (Fig. 1d) demonstrates the affinity of STAT5 to these sites. In the absence of *Stat5a*, the two copies of *Stat5b* are not sufficient to detect Stat5 binding in mammary tissue at sites E1 and E3. Similarly, no Stat5 binding was detected in T_reg_ cells in the presence of only two copies of *Stat5a*. These data support the concept that a specific threshold of STAT5 is needed for the activation of *Lrrc32*.

Our in-depth analysis revealed a highly complex *Lrrc32* super-enhancer with at least three individual STAT5-anchored enhancers, one shared between T_reg_ and mammary cells, one restricted to mammary epithelium and one specific to T_reg_ cells. STAT5 binding at a GAS motif in the enhancer shared between T_reg_ and mammary cells is structurally distinct from the site proposed by Nasrallah^4^. The binding of GR, NFIB, MED1 and Pol II to the two juxtaposed enhancers in mammary tissue suggests the presence of a multiplex super-enhancer with H3K27ac peaks spanning more than 12.5kb. Similarly, the two enhancers identified in T_reg_ cells constitute a super-enhancer.

The establishment and function of T_reg_ cells is dependent on both STAT5^9^ and GR^17^, and we propose that the enhancer-bound GR contributes the regulation of *Trrc32*. Based on mice expressing only one out of the four *Stat5* alleles, we propose that STAT5 serves as an anchor to build larger enhancer complexes. Further experiments are needed to identify the significance of the sequence identified by Nasrallah and colleagues^4^ and to establish the function of the other two candidate enhancers identified in our study. It also remains to be determined whether the risk variant rs11236797 abrogates the establishment of a specific enhancer complex. Our work also highlights the need for high-quality and deep-sequenced ChIP-seq data to obtain highly accurate information on the binding position of regulatory proteins to the genome. It remains an enigma why STAT5A/B executes opposite functions in different cell types, repression of *Lrrc32* in mammary tissue and activator in T_reg_ cells.

## Methods

### Data availability

All data were obtained from or uploaded to Gene Expression Omnibus (GEO) and Sequence Read Archive (SRA). ChIP-seq data from mammary tissue and T_reg_ cells were obtained under GSE115370, GSE145193, GSE127144, GSE40930, GSE77656 and DRA005202. RNA-seq data from mammary tissue and T_reg_ cells from WT and *Stat5* mutant mice were downloaded from GSE37646, GSE77656 and GSE128198.

## Acknowledgements

This work was supported by the Intramural Research Programs (IRPs) of National Institute of Diabetes and Digestive and Kidney Diseases (NIDDK) and utilized the computational resources of the NIH HPC Biowulf cluster (http://hpc.nih.gov).

## Author contributions

H.K.L. and L.H. designed the study. H.K.L. performed computational analysis. All authors wrote the manuscript and approved the final version.

